# EEMtoolbox: A user-friendly R package for flexible ensemble ecosystem modeling

**DOI:** 10.1101/2024.11.03.621788

**Authors:** Luz Valerie Pascal, Sarah A. Vollert, Malyon D. Bimler, Christopher M. Baker, Maude Vernet, Stefano Canessa, Christopher Drovandi, Matthew P. Adams

## Abstract

1. Forecasting ecosystem changes due to disturbances or conservation interventions is essential to improve ecosystem management and anticipate unintended consequences of conservation decisions. Mathematical models allow practitioners to understand the potential effects and unintended consequences via simulation. However, calibrating these models is often challenging due to a paucity of appropriate ecological data.
2. Ensemble ecosystem modelling (EEM) is a quantitative method used to parameterize models from theoretical ecosystem features rather than data. Two approaches have been considered to find parameter values satisfying those features: a standard accept-reject algorithm, appropriate for small ecosystem networks; and a sequential Monte Carlo (SMC) algorithm, that is more computationally efficient for larger networks. In practice, using SMC for EEM generation requires advanced statistical and mathematical knowledge, as well as strong programming skills, which might limit its uptake. In addition, current EEM approaches have been developed for only one model structure (generalized Lotka-Volterra).
3. To facilitate the usage of EEM methods we introduce EEMtoolbox, an R package for calibrating quantitative ecosystem models. Our package allows the generation of parameter sets satisfying ecosystem features, by using either the standard accept-reject algorithm or the novel SMC procedure. Our package extends the existing EEM methodology, originally developed for the generalized Lotka-Volterra model, to two additional model structures (the multi-species Gompertz, and the Bimler-Baker model), and additionally allows users to define their own model structures.
4. We demonstrate the usage of EEMtoolbox by modelling the introduction of sihek (extinct-in-the-wild) on Palmyra Atoll in the Pacific Ocean. With its simple interface, our package facilitates straightforward generation of EEM parameter sets, thus unlocks advanced statistical methods supporting conservation decisions using ecosystem network models.

## 1 Introduction

The current biodiversity crisis requires urgent interventions to preserve ecosystems (Barnosky et al., 2011; Bergstrom et al., 2021). However, conservation decisions are risky, as they can lead to unexpected negative outcomes (Roemer et al., 2002; Bergstrom et al., 2009; Buckley and Han, 2014). Our ability to anticipate consequences of interventions on ecosystems is critical to making informed decisions. Quantitative models can assist make conservation decisions by simulating effects of interventions (or disturbances) on ecosystem networks and predicting ecosystem changes (Adams et al., 2020). In practice, using these quantitative models requires large datasets to estimate driving parameters (Adams et al., 2020; Botelho et al., 2024), a difficult task in data-poor contexts such as biodiversity conservation which limits quantitative model applications (McDonald-Madden et al., 2010; Cook et al., 2013; Christie et al., 2021).

When historical data is unavailable, ensemble ecosystem modelling (EEM, Baker et al. (2017)) is a quantitative method for parameterising population models, that uses theoretical ecosystem features rather than data. EEM can therefore be applied to any ecosystem for which a network structure (or food web) has been proposed. Applications of EEM include risk analysis for the reintroduction of species (Baker et al., 2017; Peterson et al., 2021), and the management of invasive species (Rendall et al., 2021). To date, two algorithms are designed for EEM generation: an accept-reject algorithm in Baker et al. (2017), and an adaptation of the sequential Monte Carlo approximate Bayesian computation (SMC-ABC) algorithm of Drovandi and Pettitt (2011) in Vollert et al. (2024b). While the accept-reject algorithm from Baker et al. (2017) is relatively straightforward to implement, this algorithm suffers from computational limitations for moderate to large ecosystems (Peterson and Bode, 2021), limiting its application to small ecosystem networks only. In contrast, the SMC-ABC algorithm of Vollert et al. (2024b) can generate parameter sets matching the desired features orders of magnitude faster for larger and more complex ecosystem networks (Vollert et al., 2024b) but requires advanced statistical theory, mathematical knowledge and programming skills to implement. In addition, current implementations of EEM are limited to one model structure (generalised Lotka-Volterra), while other models of ecosystem dynamics are proposed in the literature to represent different types of species interactions (Gomatam, 1974; Ives et al., 2003; Bimler et al., 2024).

In this manuscript, we present a new R-package, EEMtoolbox. Our package facilitates the usage of the recent statistical advances introduced in Vollert et al. (2024b), offers a range of user-friendly functions allowing the parameter generation for three different ecosystem network model structures, and can accommodate user-defined models. The package generates an ensemble of ecosystem models using a network, and can also be used to forecast temporal changes in species abundances. As a case study, we demonstrate the usage of EEMtoolbox on the introduction of the avian top predator sihek (extinct-in-the-wild) on Palmyra Atoll, a United States minor outlying Island in the Pacific Ocean. EEMtoolbox is publicly available on Git-hub at https://anonymous.4open.science/r/EEMtoolbox-submission.

## 2 Methods

### 2.1 Modelling dynamics of ecosystem networks

Ecosystems of interacting species are commonly represented as an ecosystem network (Pimm et al., 1991; Cohen et al., 2012): a graphical representation of how populations of species or groups of species (nodes) interact (edges) with other populations. Ecosystem networks can represent predator-prey (Post et al., 2000), competitive or mutualist (Waser and Ollerton, 2006) relationships between species. Quantitative species models use this ecosystem network structure to forecast trajectories of species abundances (Murray, 2002; Ives et al., 2003; Adams et al., 2020; Baker et al., 2017). Our package EEMtoolbox can apply EEM (see Section 2.2) to three ordinary differential equation models of ecosystem networks (summarized in Table 1 and detailed on a simple predator-prey example in Appendix B), and can also accommodate user-customised models (see Appendix C).

**Table 1:**
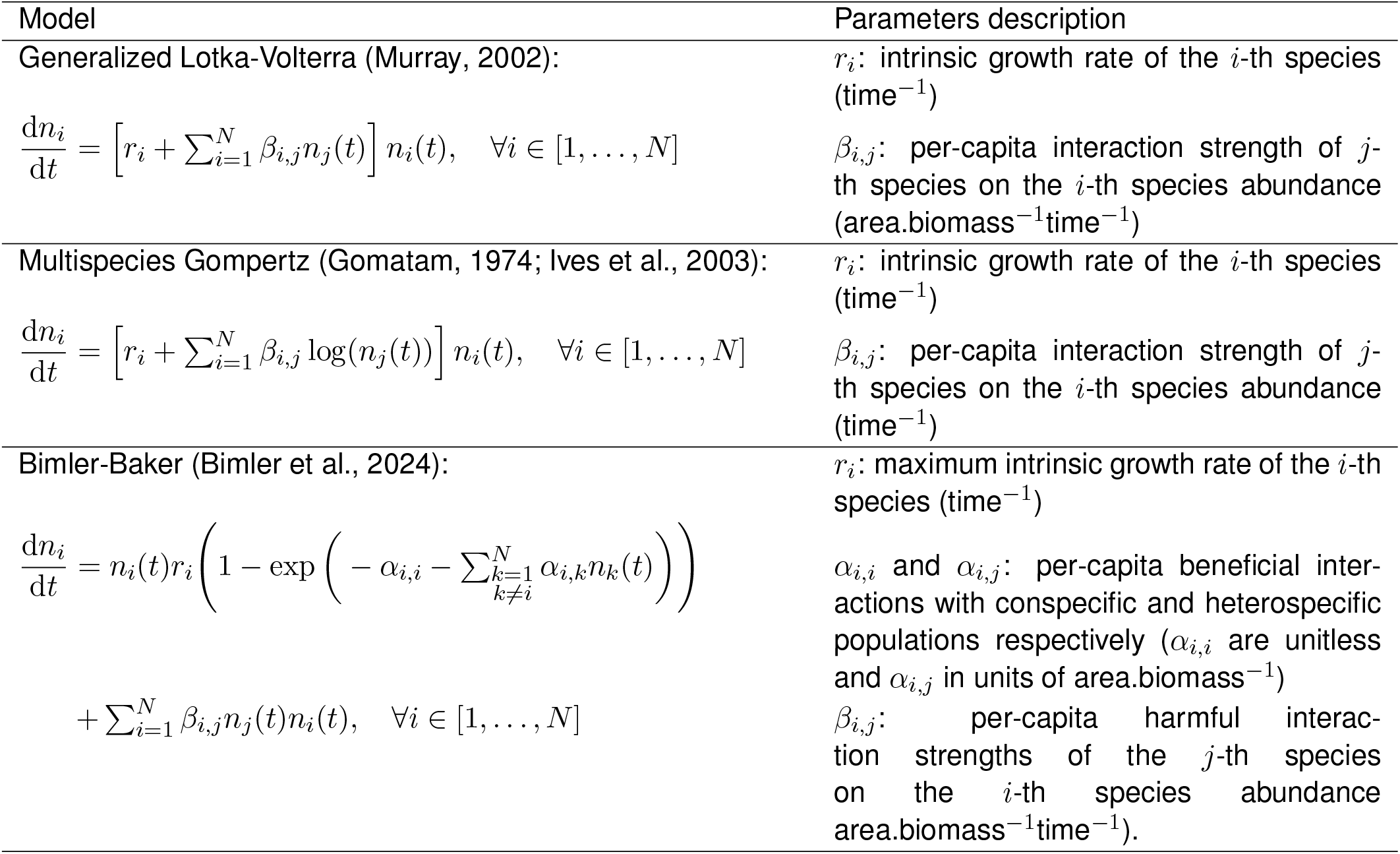
Mathematical models of ecosystem networks supported by our EEMtoolbox R-package. For all models, *n*_*i*_(*t*) represents abundance of the *i*-th species at time *t*, and *N* is the total number of species.

### 2.2 Ensemble ecosystem modelling

Ensemble ecosystem modelling (EEM) generates parameter sets using theoretical ecosystem features: feasibility (species coexistence) and stability (recovery of equilibrium populations following pulse perturbations) (Baker et al., 2017). Mathematically, feasibility implies that a steady state of the model exists in which all species’ abundances are positive, and stability implies that the steady state is Lyapunov stable (see Appendix A). Our package EEMtoolbox can apply EEM to other ecosystem models than the generalized Lotka-Volterra model using two sampling algorithms: an accept reject approach developed by Baker et al. (2017), and a sequential Monte Carlo approximate Bayesian computation algorithm (SMC-ABC, Drovandi and Pettitt (2011)) developed by Vollert et al. (2024b). Our package extends these two sampling approaches that were originally designed for the Lotka-Volterra model to other ecosystem models (see Table 1).

The EEM algorithm from Baker et al. (2017) samples parameter values of the generalised Lotka-Volterra equations, and uses an accept-reject procedure to select parameter sets that yield feasible and stable ecosystems. The process (hereafter called standard-EEM) continues until the desired number of parameter sets is generated. However, the probability of randomly generating parameter sets that meet the feasibility and stability criteria decreases dramatically as the size and complexity of the ecosystem increases (May, 1972; Allesina and Tang, 2015; Peterson et al., 2021). Thus, the usage of standard-EEM becomes practically impossible for large ecosystem networks.

The SMC-ABC algorithm from Vollert et al. (2024b) (hereafter called SMC-EEM) can generate feasible and stable parameter ensembles for larger ecosystem networks than the standard EEM approach. SMC-EEM learns from the trialled parameter sets how to better suggest parameter sets more likely to satisfy the conditions of feasibility and stability. Essentially, this approach samples parameter values, measures how poorly the sampled parameters satisfy the feasibility and stability constraints (this is called the discrepancy), and resamples and perturbs parameters to sequentially minimise the discrepancy. The target distribution of standard-EEM and SMC-EEM are identical. Thus for a large number of samples, the empirical distributions of the parameter sets produced by standard-EEM and SMC-EEM are equivalent (Vollert et al., 2024b) (see Appendix B.5). Figure 1 illustrates the difference between these approaches.

**Figure 1:**
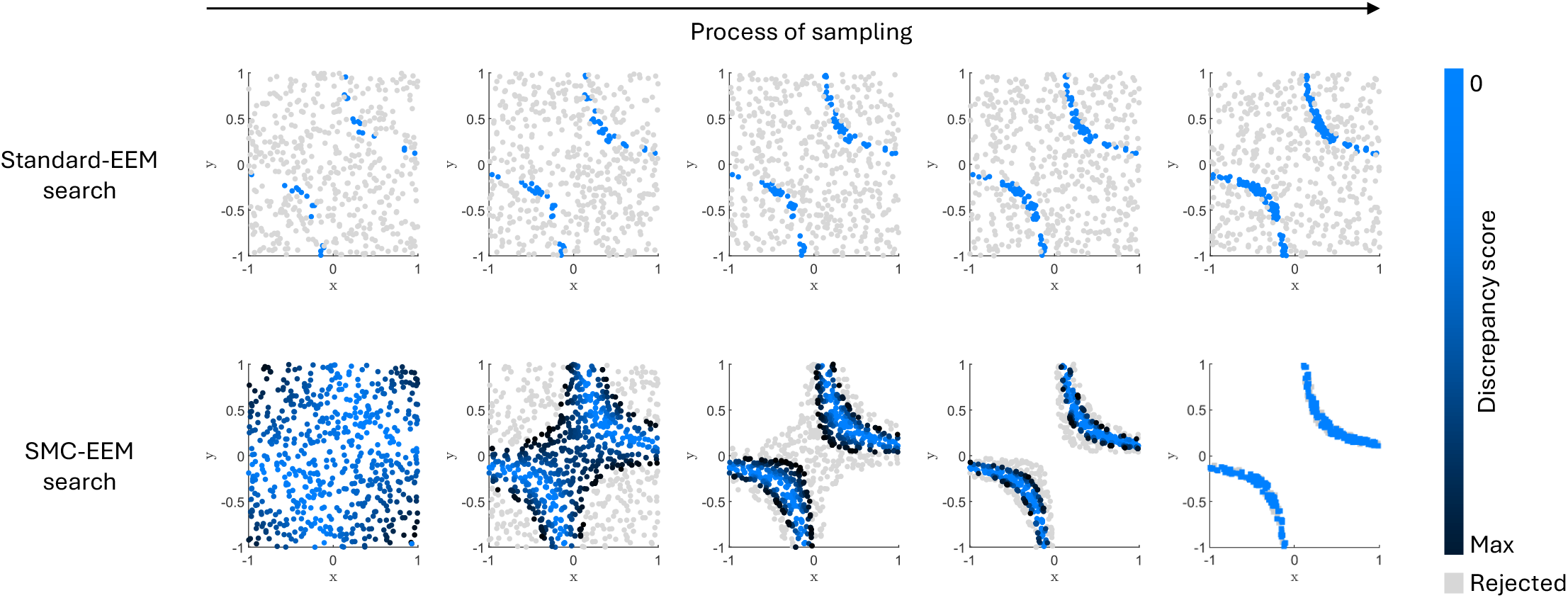
A conceptual diagram comparing the process of sampling using standard-EEM (top row) and SMC-EEM (bottom row) algorithms. The standard-EEM search method (top row) randomly samples parameters sets from a pre-specified region, rejecting parameter sets that do not meet the constraints (grey dots), and accepting those that do (blue dots). The SMC-EEM search method (bottom row) creates a discrepancy score that measures how different the desired and sampled system properties are, to sequentially adapt sampling towards the objective region. Parameter sets that meet the constraints are accepted (blue dots), while parameter sets that do not meet the constraints are rejected depending on their discrepancy score (grey dots are rejected and dark blue dots will be resampled from and perturbed to sequentially minimise the discrepancy score). This two dimensional illustration aims to obtain parameter sets where 0.1 ≤ *xy* ≤ 0.15 (curves in top right and bottom left region of parameter space); however, for EEM these algorithms aim to obtain regions of parameter space that yield feasible and stable ecosystem models.

## 3 Demonstrating the functionality of the EEMtoolbox R-package

### 3.1 Sihek case study

Sihek, a charismatic bird species, endemic to Guam in the North-Western Pacific, went extinct in the wild following the accidental introduction of the invasive brown tree snake (*Boiga irregularis*) in its native range (Savidge, 1986; Fish et al., 1990). Re-establishing a population of sihek in the wild is urgent due to the limited capacity of zoos and the increasing genetic concerns related to the species’ small current population size (Trask et al., 2021). Since reintroducing sihek to their endemic range is not yet feasible, a conservation introduction to Palmyra Atoll is being considered in the meantime. Due to their ecology, Canessa et al. (2022) anticipate sihek replacing carnivorous crabs and cane spiders as top predators in Palmyra Atoll upon introduction.

We use EEM to explore the potential impacts of sihek introduction on the abundances of the resident species of Palmyra Atoll. Following Canessa et al. (2022), we model changes in the abundance of all species predicted to interact directly with sihek (nine species or groups of species; Figure 2) from before the introduction up to 10 years post-introduction. We use the upper bounds for the species’ growth rates and types of interaction derived by Canessa et al. (2022), summarized in Table S1 and Figure 2.

**Figure 2:**
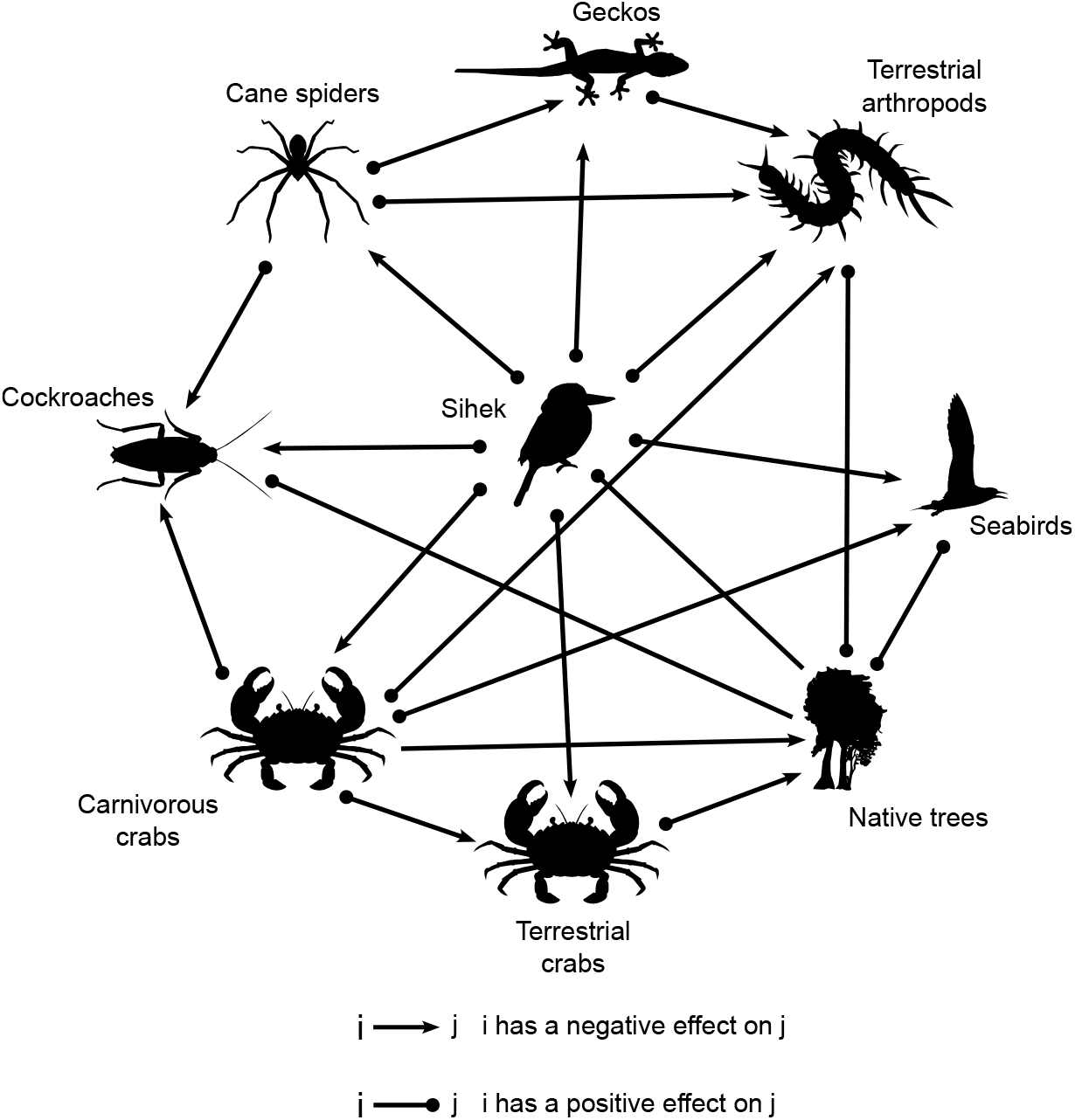
Species interactions network for the sihek case study. All species also possess a negative intraspecies interaction, as is commonly assumed in ecosystem network models (Baker et al., 2017; Adams et al., 2020)

### 3.2 Generating ensembles of ecosystem models

`EEM` is the main function of the R-package and it can generate an ensemble of parameter sets that yield feasible and stable ecosystem networks (see Table 2 for relevant arguments of the function). Usage of the function for a 2-species example is provided in Appendix B.2. A second more illustrative example is provided by the following code which generates 5,000 parameter sets satisfying the feasibility and stability features of the whole ecosystem, for the generalized Lotka-Volterra model of the sihek case-study (see Table S1):

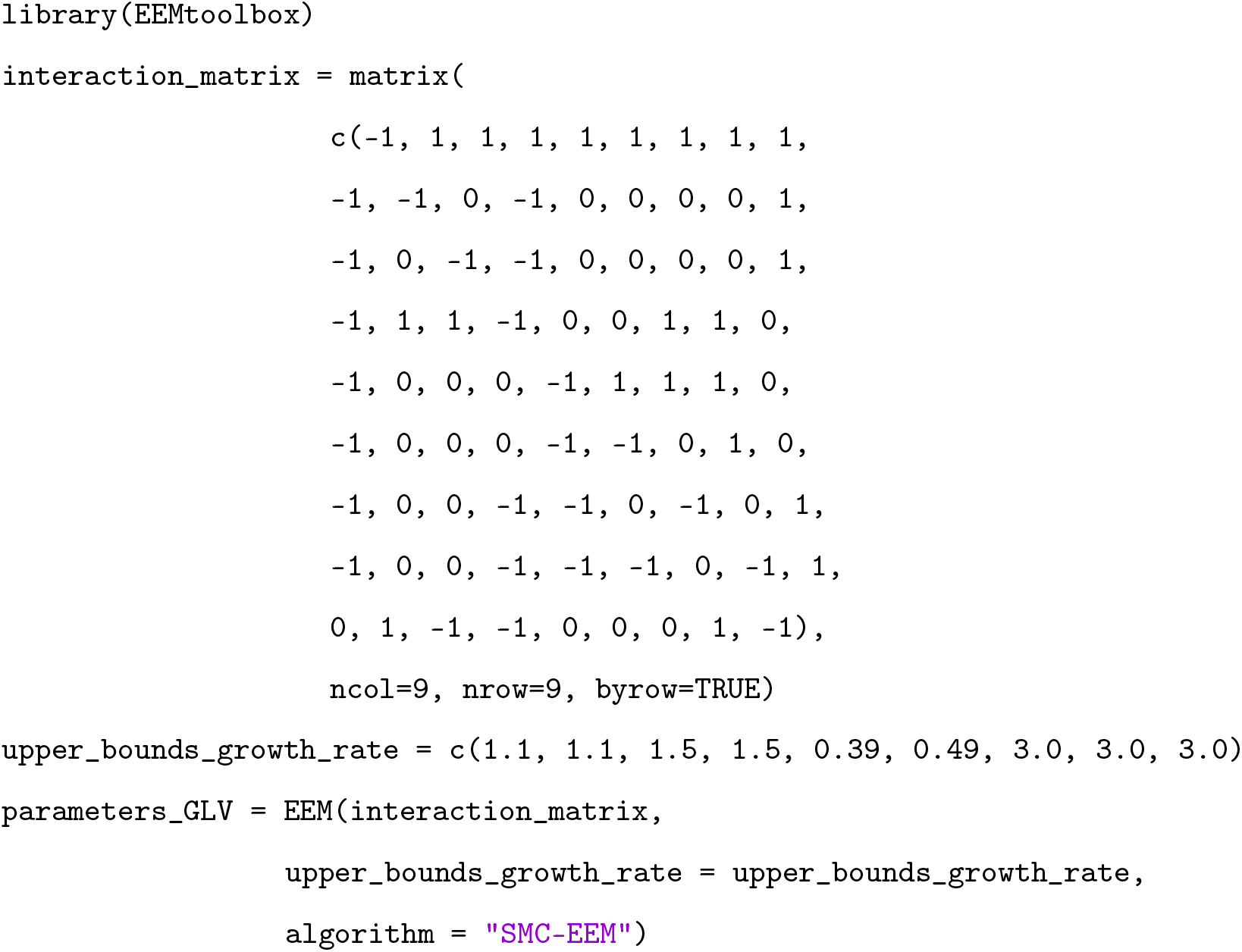

**Table 2:** Summary of relevant arguments of the `EEM` function. For the `model` argument, “GLV” stands for generalized Lotka-Volterra. The argument `n_cores` sets the number of clusters used for generating parameter sets. The default value `1L` signifies that the sampling process is sequential. Setting `n_cores` to a value larger than `1L` parallelises the sampling process on the number of desired clusters. We refer the reader to the documentation of the function for further details including possible parameter values for each argument.

| Argument name | Description | Default value |
| --- | --- | --- |
| `interaction_matrix` | Bounds of interaction matrix for the ecosystem network | - |
| `lower_bounds_growth_rate` | Lower bounds for sampled growth rates ($r_i$) | 0 |
| `upper_bounds_growth_rate` | Upper bounds for sampled growth rates ($r_i$) | 5 |
| `n_ensemble` | Number of desired parameter sets | 5,000 |
| `model` | Model representing species interactions | "GLV" |
| `algorithm` | Sampling algorithm | "standard-EEM" |
| `n_cores` | Number of cores used for sampling. | 1L |

This code (using the SMC-EEM algorithm) allowed us to generate 5,000 parameter sets in approximately 4 minutes on a virtual machine (AMD EPYC 7702 64-Core Processor with 64 GB of RAM, 16 virtual processors, and using 15 of the 16 available threads). In contrast, generating 5,000 parameter sets using the standard-EEM algorithm was achieved in approximately 2 days and 7 hours on the same virtual machine.

### 3.3 Generating projections of future abundances

The function `plot_projections` uses the obtained parameter sets to forecast changes in species abundances, by numerically solving ordinary differential equations (ODEs, see Algorithm S1). This function inputs parameter sets (e.g. generated by the `EEM` function), initial species abundances, and a time window for the forecast. For each parameter set, `plot_projections` solves the corresponding ODE using the function `ode` from the R-package `deSolve` (Soetaert et al., 2010). Finally, the function estimates the median abundance and 95% prediction intervals for each species and at each time step, then plots the corresponding graph (see Figure 3). We summarize in Table 3 all relevant parameters of the function. The following code forecasts species abundances over 10 years, when the initial abundance of sihek is 0.1 (individuals per hectare), and the other species are 1 (individuals per hectare). For a more detailed demonstration of this function on a simple two-species example see Appendix B.3, and Appendix B.4 for projections scaled to the steady state.

**Table 3:** Summary of parameters for the `plot_projections` function. All these parameters require user specification.

| Argument name | Description |
| --- | --- |
| `parameters` | List like object of ensemble of parameters (outputs of EEM) |
| `model` | Model representing species interactions (default "GLV") |
| `initial_condition` | Initial species abundances |
| `t_window` | Time window for projections |

**Figure 3:**
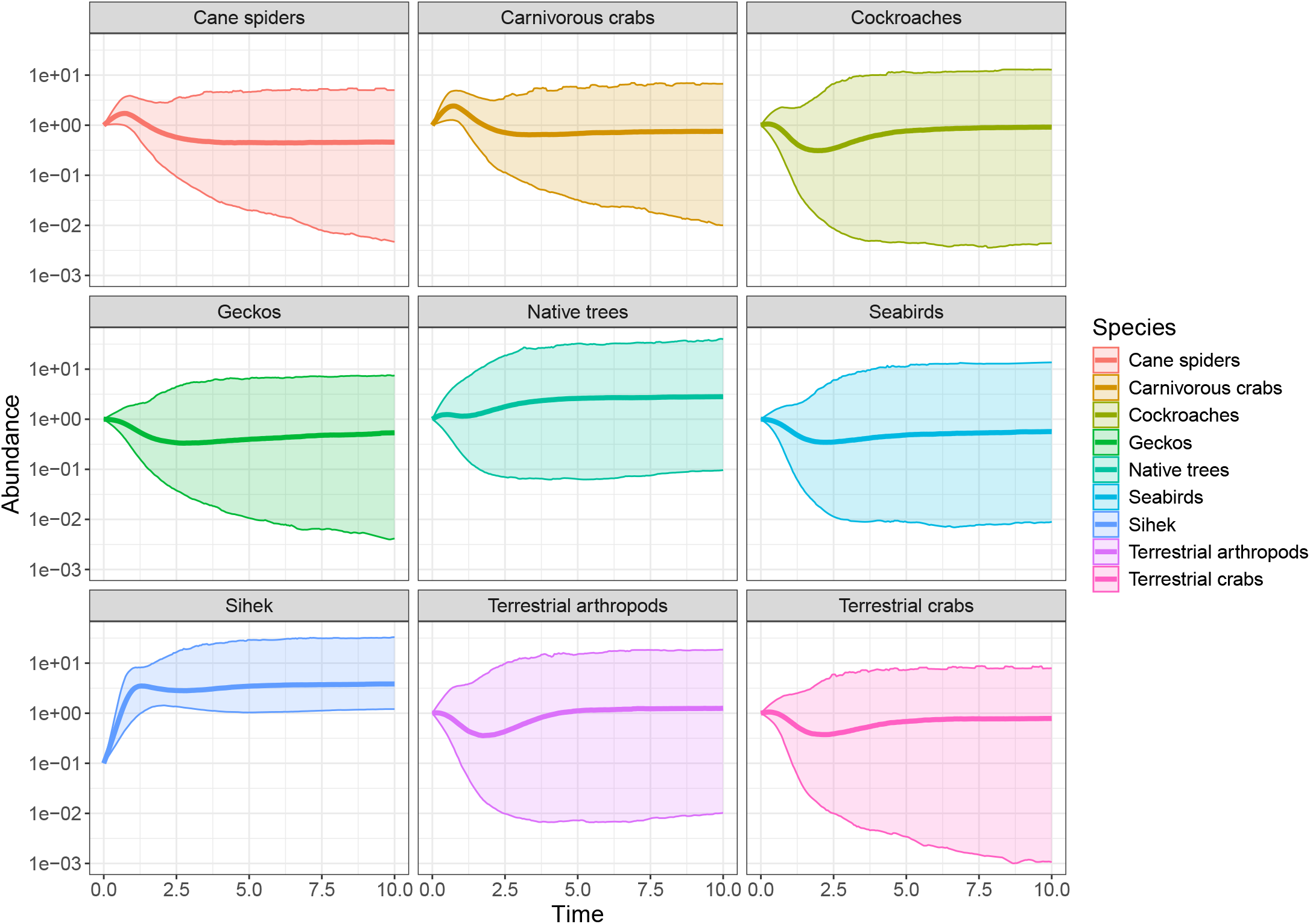
Forecasts of species abundances over 10 years for the sihek case study using the generalized Lotka-Volterra model. For illustrative purposes, we set the initial abundance of sihek to 0.1 (individuals per hectare), and the other species are 1 (individuals per hectare). Thicker lines represent the mean abundance and thin lines the 95% prediction interval. Forecasts for the multi-species Gompertz and the Bimler-Baker model are provided in Appendix E (Fig. S5 and S6)

~~~
plot_projections(parameters=parameters_GLV,
              initial_condition=c(0.1, 1, 1, 1, 1, 1, 1, 1, 1),
              t_window = c(0,10))
~~~

## 4 Discussion

This paper showcases a new R-package EEMtoolbox that quickly and easily generates parameter sets for ecosystem model ensembles, for up to three different model structures, as well as for user defined models. The package can either sample parameter values using a standard accept-reject approach recommended for small networks as in Baker et al. (2017), or via an SMC-ABC algorithm recommended for larger networks (Drovandi and Pettitt, 2011; Vollert et al., 2024b). By facilitating the usage of these advanced statistical methods within an R-package, EEM can now be applied to parameterize models of a broad range of ecosystem network structures, yielding immediate utility for improving conservation decisions based on these models. Our package extends EEM to three model structures (generalized Lotka-Volterra, the multi-species Gompertz and the Bimler-Baker models), and can be applied to user defined model structures (see Appendix C).

We acknowledge other factors might hinder EEM applications. First, EEM assumes the studied ecosystem will reach a stable and feasible equilibrium, which is convenient but not always appropriate (Cuddington, 2001; Francis et al., 2021; Vollert et al., 2024a). Second, EEM requires defining a species interaction network, which remains a difficult task (see for example Peterson et al. 2021). Uncertainties in the ecosystem network (such as existence or not of direct interactions between species, and misspecification of interaction) might result in misleading predictions of species abundances (Adams et al., 2020). Third, EEM can provide new information but not direct recommendations for ecosystem management (Baker et al., 2017). In our case study, the sihek Recovery Team used EEM as an additional source of insights about predation and competition, following a broad risk screening for sihek introduction to Palmyra Atoll (Vernet et al., 2024). EEM is therefore especially useful when rapid exploration of scenarios and frequent updating of projections and risk assessments are desired by decision-makers (Canessa et al., 2022).

## Acknowledgments

We thank the QUT Centre for Data Science for funding this research through the Second Byte Funding Program 2024. SAV and MPA acknowledge funding from an Australian Research Council Discovery Early Career Researcher Award (DE200100683). LVP acknowledges funding from an Australian Research Council Discovery Early Career Researcher Award (DE200101791). LVP and CD acknowledge funding from an Australian Research Council Discovery Project (DP200102101). MDB acknowledges funding from the Botany Foundation. Funding for the sihek case study was provided through Guam Department of Agriculture (DAWR) and U.S. Fish and Wildlife Service Endangered Species Section 6 grant (Federal Award F20AF12070). LVP and MPA acknowledge funding from the ARC SRIEAS Grant SR200100005 Securing Antarctica’s Environmental Future. We thank Oakes Holland for helpful discussions during the development of this manuscript. We thank Suzanne Medina, Megan Laut and John Ewen (Sihek Recovery Team) for support and advice.

## Data availability

The R-package is available at: https://anonymous.4open.science/r/EEMtoolbox-submission.

## Authors contributions

L.P. led the research, developed the R-package, and led the writing. M.A. conceived the project and supervised the study. S.V. contributed to the idea and methodology design. M.B. and C.B. designed and provided insights towards the model development. M.V. and S.C. provided the case study. M.B., M.V. and S.C. tested the R-package. C.D. revised the manuscript, and provided insights on the statistical methods used. All authors edited the manuscript, contributed to interpretation of the results, and gave final approval for publication.

