## Supplemental material for "EEMtoolbox: A user-friendly R package for flexible ensemble ecosystem modeling"

#### A Feasibility and stability criteria

In Section 2.2, we stated that feasibility and stability criteria for each of the three models included in the EEMtoolbox package (generalised Lotka-Volterra, multi-species Gompertz and Bimler-Baker models), can be described mathematically by the requirement of (1) a steady state existing in which all species abundances are positive (feasibility) and (2) that this steady state is Lyapunov stable (stability). Here, we derive the mathematical conditions of feasibility and stability for all three models included in the EEMtoolbox package.

##### A.1 Generalized Lotka-Volterra

These equations can be equivalently expressed in a vector form as:

$$\frac{\mathrm{d}\tilde{n}}{\mathrm{d}t} = [\tilde{r} + \mathbf{A}\tilde{n}] \circ \tilde{n},$$

where $\tilde{n}$ is the vector of abundances, $\tilde{r}$ is the vector of intrinsic growth rates, $\mathbf{A}$ is the $N \times N$ interaction matrix of per-capita interaction strengths where $A_{i,j} = \beta_{i,j}$, and $\circ$ is the Hadamard (element wise) product.

###### A.1.1 Feasibility

The feasibility conditions require finding a steady state of the model that that is positive for all species. To find the steady-state of the generalized Lotka-Volterra model, we solve for $\tilde{n}^*$:

$$\frac{\mathrm{d}\tilde{n}^*}{\mathrm{d}t} = [\tilde{r} + \mathbf{A}\tilde{n}^*] \circ \tilde{n}^* = 0.$$

The solution to this equation is:

$$\tilde{n}^* = -\mathbf{A}^{-1}\tilde{r}.$$

The feasibility condition is satisfied if all the elements of $\tilde{n}^*$ are strictly positive i.e. $\tilde{n}^* = -\mathbf{A}^{-1}\tilde{r} > 0$.

##### A.1.2 Stability

To verify the (Lyapunov) stability of the solution $\mathbf{n}^*$, we need to calculate the eigenvalues of the Jacobian matrix $J$ at equilibrium $\mathbf{n}^*$. All the elements $J_{i,j}$ of the Jacobian matrix $J$ are defined as:

$$J_{i,j} = \left(\frac{\partial f_i}{\partial n_j}\right)_{\tilde{n}=\tilde{n}^*}, \quad \text{where } f_i(\tilde{n}) = \left[r_i + \sum_{i=1}^{N} \beta_{i,j} n_j(t)\right] n_i(t).$$

These elements for the generalized Lotka-Volterra model are thus defined as:

$$\begin{cases} J_{i,j} = \beta_{i,j} n_i^*, \quad \forall i \neq j \\ J_{i,i} = \underbrace{r_i + \sum_{i=1}^{N} \beta_{i,j} n_j^*(t)}_{=0 \text{ at equilibrium}} + \beta_{i,i} n_i^*(t) = \beta_{i,i} n_i^*. \end{cases}$$

Therefore, $J_{i,j} = \beta_{i,j} n_i^*, \quad \forall i, j.$

The system is considered stable if the real part of all eigenvalues ($\lambda_i$) of the Jacobian matrix are negative, i.e. $Re(\lambda_i) \leq 0, \quad \forall i = 1, \cdots, N.$

#### A.2 Gompertz

##### A.2.1 Feasibility

The steady states of the Gompertz model ODEs are given by:

$$\tilde{n}^* = \exp(-\mathbf{A}^{-1}\tilde{r}),$$

where $\exp$ is taken element-wise.

This solution $\tilde{n}^*$ is always positive, because of the exponential function, so the ecosystem is always feasible in the multi-species Gompertz model. The reader will note here that the steady states of the multi-species Gompertz model are the exponent of the steady states for the generalized Lotka-Volterra.

##### A.2.2 Stability

By following the same procedure as for the generalized Lotka-Volterra model, all elements for the Jacobian matrix are given by:

$$J_{i,j} = \beta_{i,j}\frac{n_i^*}{n_j^*}.$$

The system is considered stable if the real part of all eigenvalues ($\lambda_i$) of the Jacobian matrix are negative, i.e. $Re(\lambda_i) \leq 0, \quad \forall i = 1, \cdots, N.$

#### A.3 Bimler-Baker

##### A.3.1 Feasibility

The steady states of the Bimler-Baker model are given by:

$$\frac{\mathrm{d}n_i^*}{\mathrm{d}t} = r_i n_i^* \left[1 - e^{-\alpha_{i,i} - \sum_{k\neq i}\alpha_{i,k}n_k^*}\right] + n_i^* \sum_{i=1}^{N} \beta_{i,j} n_j^* = 0.$$

Under the assumption that $n_i^* \neq 0$, we obtain:

$$r_i \left[1 - e^{-\alpha_{i,i} - \sum_{k\neq i}\alpha_{i,k}n_k^*}\right] + \sum_{i=1}^{N} \beta_{i,j} n_j^* = 0.$$

Because this set of equations is non-linear, it is solved numerically, for example using Newton's method (Meza, 2011).

##### A.3.2 Stability

To verify the (Lyapunov) stability of the solution $\mathbf{n}^*$, we need to calculate the eigenvalues of the Jacobian matrix $J$ at equilibrium $\mathbf{n}^*$. The Jacobian matrix $J$ is defined as:

$$J_{i,j} = \left(\frac{\partial f_i}{\partial n_j}\right)_{n=n^*}, \quad \text{where } f_i(\tilde{n}) = r_i n_i(t)\left[1 - e^{-\alpha_{i,i} - \sum_{k\neq i}\alpha_{i,k}n_k(t)}\right] + n_i(t)\sum_{i=1}^{N}\beta_{i,j}n_j(t).$$

The Jacobian is thus defined as:

$$J_{i,i} = \beta_{i,i} n_i^*,$$
$$J_{i,j} = r_i n_i^* \alpha_{i,j} \exp(-\alpha_{i,i} - \sum_{k\neq i}\alpha_{i,k}n_k^*) + \beta_{i,j}n_i^*, \quad \forall i \neq j.$$

#### B Two-species ecosystem example: foxes and rabbits

In this section, we illustrate the main functionalities of our package using a simple example. We consider an ecosystem with two interacting species: a prey (rabbits) and a predator (foxes), see Fig. S1.

##### B.1 Model definition

We assume foxes have a negative impact on the rabbits, and foxes have a negative interaction with other foxes (intraspecific competition). Rabbits have a positive impact on foxes (provide food) and a negative impact on other rabbits (intraspecific competition). The interaction matrix ($\beta$) for this fox-rabbit example has the following structure:

$$\begin{array}{c} \\ \text{Fox} \\ \text{Rabbit} \end{array} \begin{array}{c} \begin{array}{cc} \text{Fox} & \text{Rabbit} \end{array} \\ \begin{pmatrix} \beta_{\mathrm{F,F}} & \beta_{\mathrm{F,R}} \\ \beta_{\mathrm{R,F}} & \beta_{\mathrm{R,R}} \end{pmatrix} \end{array} = \begin{array}{c} \\ \text{Fox} \\ \text{Rabbit} \end{array} \begin{array}{c} \begin{array}{cc} \text{Fox} & \text{Rabbit} \end{array} \\ \begin{pmatrix} -b_{\mathrm{F,F}} & +b_{\mathrm{F,R}} \\ -b_{\mathrm{R,F}} & -b_{\mathrm{R,R}} \end{pmatrix} \end{array}, \quad (1)$$

where $b_{i,j} = |\beta_{i,j}|$. The system of generalized Lotka-Volterra equations describing this ecosystem is then:

$$\begin{aligned} \frac{\mathrm{d}n_{\mathrm{F}}}{\mathrm{d}t} &= \Big[r_{\mathrm{F}} - b_{\mathrm{F,F}} n_{\mathrm{F}}(t) + b_{\mathrm{F,R}} n_{\mathrm{R}}(t)\Big] n_{\mathrm{F}}(t), \\ \frac{\mathrm{d}n_{\mathrm{R}}}{\mathrm{d}t} &= \Big[r_{\mathrm{R}} - b_{\mathrm{R,F}} n_{\mathrm{F}}(t) - b_{\mathrm{R,R}} n_{\mathrm{R}}(t)\Big] n_{\mathrm{R}}(t). \end{aligned}$$

Similarly, the system of equations for the multi-species Gompertz model is:

$$\begin{aligned} \frac{\mathrm{d}n_{\mathrm{F}}}{\mathrm{d}t} &= \Big[r_{\mathrm{F}} - b_{\mathrm{F,F}} \log\big(n_{\mathrm{F}}(t)\big) + b_{\mathrm{F,R}} \log\big(n_{\mathrm{R}}(t)\big)\Big] n_{\mathrm{F}}(t), \\ \frac{\mathrm{d}n_{\mathrm{R}}}{\mathrm{d}t} &= \Big[r_{\mathrm{R}} - b_{\mathrm{R,F}} \log\big(n_{\mathrm{F}}(t)\big) - b_{\mathrm{R,R}} \log\big(n_{\mathrm{R}}(t)\big)\Big] n_{\mathrm{R}}(t). \end{aligned}$$

As opposed to the generalized Lotka-Volterra and the multi-species Gompertz models that use one interaction matrix (Eq. 1), the Bimler-Baker model separates beneficial and harmful interactions into two interaction matrices ($\alpha$ and $\beta$ respectively). For the fox-rabbit example, these interaction matrices $\alpha$ and $\beta$ are:

$$\alpha = \begin{array}{c} \\ \text{Fox} \\ \text{Rabbit} \end{array} \begin{array}{c} \begin{array}{cc} \text{Fox} & \text{Rabbit} \end{array} \\ \begin{pmatrix} +a_{\mathrm{F,F}} & +a_{\mathrm{F,R}} \\ 0 & +a_{\mathrm{R,R}} \end{pmatrix} \end{array} \text{ and } \beta = \begin{array}{c} \\ \text{Fox} \\ \text{Rabbit} \end{array} \begin{array}{c} \begin{array}{cc} \text{Fox} & \text{Rabbit} \end{array} \\ \begin{pmatrix} -b_{\mathrm{F,F}} & 0 \\ -b_{\mathrm{R,F}} & -b_{\mathrm{R,R}} \end{pmatrix} \end{array}, \quad (2)$$

where all $a_{i,j}$ and $b_{i,j}$ are strictly positive. The system of equations for the Bimler-Baker model is then:

$$\frac{\mathrm{d}n_{\mathrm{F}}}{\mathrm{d}t} = \left[ r_{\mathrm{F}} \left( 1 - e^{-a_{\mathrm{F,F}} - a_{\mathrm{F,R}} n_{\mathrm{R}}(t)} \right) - b_{\mathrm{F,F}} n_{\mathrm{F}}(t) \right] n_{\mathrm{F}}(t),$$

$$\frac{\mathrm{d}n_{\mathrm{R}}}{\mathrm{d}t} = \left[ r_{\mathrm{R}} \left( 1 - e^{-a_{\mathrm{R,R}}} \right) - b_{\mathrm{R,F}} n_{\mathrm{F}}(t) - b_{\mathrm{R,R}} n_{\mathrm{R}}(t) \right] n_{\mathrm{R}}(t).$$

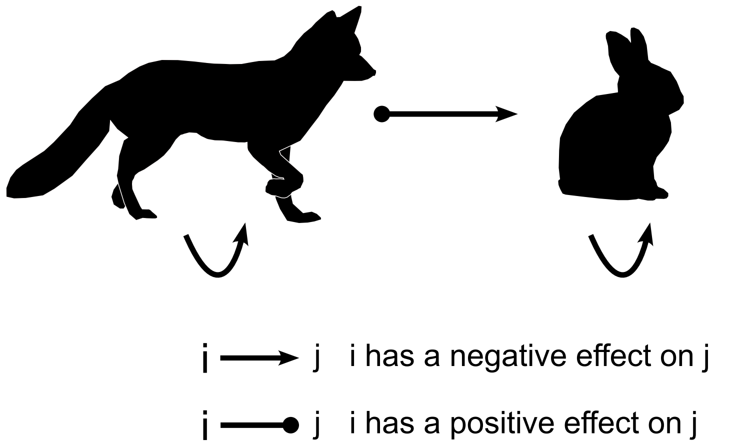


Figure S1: Ecosystem interaction network for the fox-rabbit example.

##### B.2 Generating ensembles of ecosystem models

The main function of our R-package is `EEM` and is designed to generate an ensemble of parameter sets that yield feasible and stable ecosystem networks. To sample such parameters following the generalized Lotka-Volterra model, users only need to input the bounds of an interaction matrix such as Equation (1) to generate ecosystem ensembles. The following code generates a set of parameters for the fox-rabbit example using the generalized Lotka-Volterra equations:

```
library(EEMtoolbox)
interaction_matrix = matrix(c(-1,-1,1,-1), nrow=2, ncol=2)}
parameters_GLV = EEM(interaction_matrix)
```

The first line (shown above) loads all the functions of the `EEMtoolbox` package. The second defines the bounds of the interaction matrix of the studied ecosystem through the function `matrix`. Here, the argument `c(-1,-1,1,-1)` is a vector defining the bounds (positive or negative) of parameters $\beta_{\mathrm{F,F}}$, $\beta_{\mathrm{R,F}}$, $\beta_{\mathrm{F,R}}$ and $\beta_{\mathrm{R,R}}$ respectively, as well as the upper and lower bounds for the sampling procedure. Here, these parameters $\beta$ will be sampled from the intervals $[-1,0]$, $[-1,0]$, $[0,1]$ and $[-1,0]$ respectively. If a positive (resp. negative) number ($x$) were chosen instead of 1, 0 or -1, the sampling interval would be $[0,x]$ (resp. $[x,0]$). The arguments `nrow=2, ncol=2` signify that the interaction matrix has 2 rows and 2 columns (as there are 2 interaction species in the ecosystem). The third and final line runs the EEM function with interaction_matrix as argument. Here, EEM samples 5,000 parameter sets (see Table 2 for default values of the function EEM) satisfying the feasibility and stability features of the ecosystem. These parameter sets are bounded as follows: the intrinsic growth rates ($r$) are sampled within the interval $[0, 5]$ (EEM default values), and the species interaction terms ($\beta$) are bounded within the aforementioned intervals.

Sampling parameters for the multi-species Gompertz model only requires changing the argument model in the third line as follows, since the element bounds of the interaction matrix (interaction_matrix) remain the same as for the generalized Lotka-Volterra model:

```
parameters_Gompertz = EEM(interaction_matrix, model="Gompertz")
```

To generate parameter sets using the Bimler-Baker model, the user must provide the bounds of elements of two interaction matrices (one for the beneficial interactions, and one for the detrimental interactions):

```
alphas_matrix = matrix(c(1,0,1,1), nrow=2, ncol=2)
betas_matrix = matrix(c(-1,-1,0,-1), nrow=2, ncol=2)
parameters_Baker = EEM(list(alphas_matrix, betas_matrix), model="Bimler-Baker")
```

Here, the first and second lines define the bounds of the interaction matrices (Eq. 2). In the first line, the argument c(1,0,1,1) is a vector defining the bounds of parameters $\alpha_{\mathrm{F,F}}$, $\alpha_{\mathrm{R,F}}$, $\alpha_{\mathrm{F,R}}$ and $\alpha_{\mathrm{R,R}}$ respectively. Here, these parameters $\alpha_{\mathrm{F,F}}$, $\alpha_{\mathrm{F,R}}$ and $\alpha_{\mathrm{R,R}}$ will all be sampled from the interval $[0, 1]$, and the parameter $\alpha_{\mathrm{R,F}}$ will be 0. Similarly, in the second line, the argument c(-1,-1,0,-1) defines the bounds of parameters $\beta_{\mathrm{F,F}}$, $\beta_{\mathrm{R,F}}$, $\beta_{\mathrm{F,R}}$ and $\beta_{\mathrm{R,R}}$ respectively. Here, $\beta_{\mathrm{F,F}}$, $\beta_{\mathrm{R,F}}$ and $\beta_{\mathrm{R,R}}$ will all be sampled from the interval $[-1, 0]$, and $\beta_{\mathrm{F,R}}$ will be 0. If a positive (resp. negative) number ($x$) were chosen instead of 1, 0 or -1, the sampling interval would be $[0, x]$ (resp. $[x, 0]$). The third and final line runs the EEM function with a list object as input for the argument interaction_matrix, and the model argument set to "Bimler-Baker". The two elements of the list, in this third line, are the matrices $\alpha$ and $\beta$. In all of the above codes, the sampling algorithm was not selected by the user, so was set to the default "standard-EEM" algorithm (see Table 2), which is appropriate for small networks. For larger networks, the "SMC-EEM" algorithm should be used. The manuscritp shows a code example (see Section 3.2) which utilises the "SMC-EEM" algorithm for a nine-species ecosystem.

##### B.3 Generating projections of future abundances of species

One key usage of obtaining an ensemble of feasible and stable ecosystems is to forecast ecosystem changes due to interventions or disturbances (Section 2.1). These forecasts can be achieved by solving the ordinary differential equations (ODEs) for a given parameter set, and a given initial condition (or initial abundance). The following equation represents the system of ODEs for the generalized Lotka-Volterra model, with parameters $r_{\mathrm{F}}$, $r_{\mathrm{R}}$, $\beta_{\mathrm{F,F}}$,

$\beta_{\rm F,R}$, $\beta_{\rm R,F}$, $\beta_{\rm R,R}$ (obtained using the `EEM` function) and initial abundances $n_{\rm F}^0$ for foxes and $n_{\rm R}^0$ for rabbits (provided by the user).

$$\left.\begin{aligned}\frac{\mathrm{d}n_{\rm F}}{\mathrm{d}t} &= r_{\rm F}n_{\rm F}(t) - \beta_{\rm F,F}n_{\rm F}(t)^2 + \beta_{\rm F,R}n_{\rm F}(t)n_{\rm R}(t),\\ \frac{\mathrm{d}n_{\rm R}}{\mathrm{d}t} &= r_{\rm R}n_{\rm R}(t) - \beta_{\rm R,F}n_{\rm F}(t)n_{\rm R}(t) - \beta_{\rm R,R}n_{\rm R}(t)^2\end{aligned}\right\}\text{ solved from } t=0 \text{ to } t=s$$

given that $n_{\rm F}(0) = n_{\rm F}^0, \quad n_{\rm R}(0) = n_{\rm R}^0.$

In this context, solving a system of ODEs yields predicted time-serie for how species abundances are going to change over time, given an initial abundance.

The function `plot_projections` numerically solves the ODEs for the full ensemble of parameter sets, and forecasts species abundances (Algorithm S1). This function requires input parameter sets (e.g. generated by the `EEM` function), initial species abundances (e.g. $n_{\rm F}^0, n_{\rm R}^0$), and a time window for the forecast of abundance change. For each parameter set, `plot_projections` solves the corresponding system of ODEs (see equation above) by calling the `ode` function of the R-package `deSolve` (Soetaert et al., 2010). Finally, the function estimates the median abundance at each time step, and for each species, as well as the 95% prediction intervals, and plots the corresponding graph. Table 3 summarises all relevant inputs to this function. As an example, the following code generates abundance projections over 5 years when the abundance of both species are initially set to 1 individual per hectare, for each of the three models coded in the package (Figure S2). Example code for the `plot_projections` function applied to a nine-species ecosystem is also provided in Section 3.3.

```
plot_projections(parameters_GLV, initial_condition = c(1,1), t_window = c(0,5))
plot_projections(parameters_Gompertz, model="Gompertz",
                 initial_condition = c(1,1), t_window = c(0,5))
plot_projections(parameters_Baker, model="Bimler-Baker",
                 initial_condition = c(1,1), t_window = c(0,5))
```

**Algorithm S1** Pseudo-code of the `plot_projections` function

**Input:** Parameter sets, initial species abundance ($n_{\rm F}^0, n_{\rm R}^0$), time window
**Output:** Plot of median abundance over time, with 95% prediction intervals and for each species
1: **for** For all parameter sets $i \in [1, \cdots, M]$ **do**
2: Numerically solve a system of ODEs with current parameters, initial condition $n_{\rm F}^0, n_{\rm R}^0$, between $t = 0$ and $t = T$.
3: Store the computed trajectories of species abundances $n_{\rm F}(t), n_{\rm R}(t)$
4: Using the $M$ computed trajectories $n_{\rm F}(t), n_{\rm R}(t)$, compute median $n_{\rm F}(t), n_{\rm R}(t)$, and 95% prediction intervals.
5: **return** Plot of median abundance over time, with 95% prediction intervals and for each species

Although the parameters bounds are the same for all models during the sampling procedure, the projected species' abundances are unsurprisingly different for each model. For example, for the generalized Lotka-Volterra model, the fox abundance varies between 0 and 30, with and average of $\approx 6$ and the rabbit abundance between 0 and 15, with an average of $\approx 3$. In contrast, the projected species' abundances for the multi-species Gompertz model are significantly higher than those projected with the generalized Lotka-Volterra model. This is because the steady state of the multi-species Gompertz model is equal to the exponential of the steady state for the Lotka-Volterra model, leading to higher projected abundances (see Appendix A.2). Similarly, the projected species's abundances for the Bimler-Baker model are lower than those projected by the generalized Lotka-Volterra or multi-species Gompertz models. This is due to the introduction of the exponential term in the Bimler-Baker equations (see Table 1 in main manuscript), which diminishes the effect of the intrinsic growth rate, as well as the exclusion of the positive interactions in the final term $\sum \beta_{i,j} n_i(t) n_j(t)$ of this model (in comparison to the other two models which do include the positive interactions in this term).

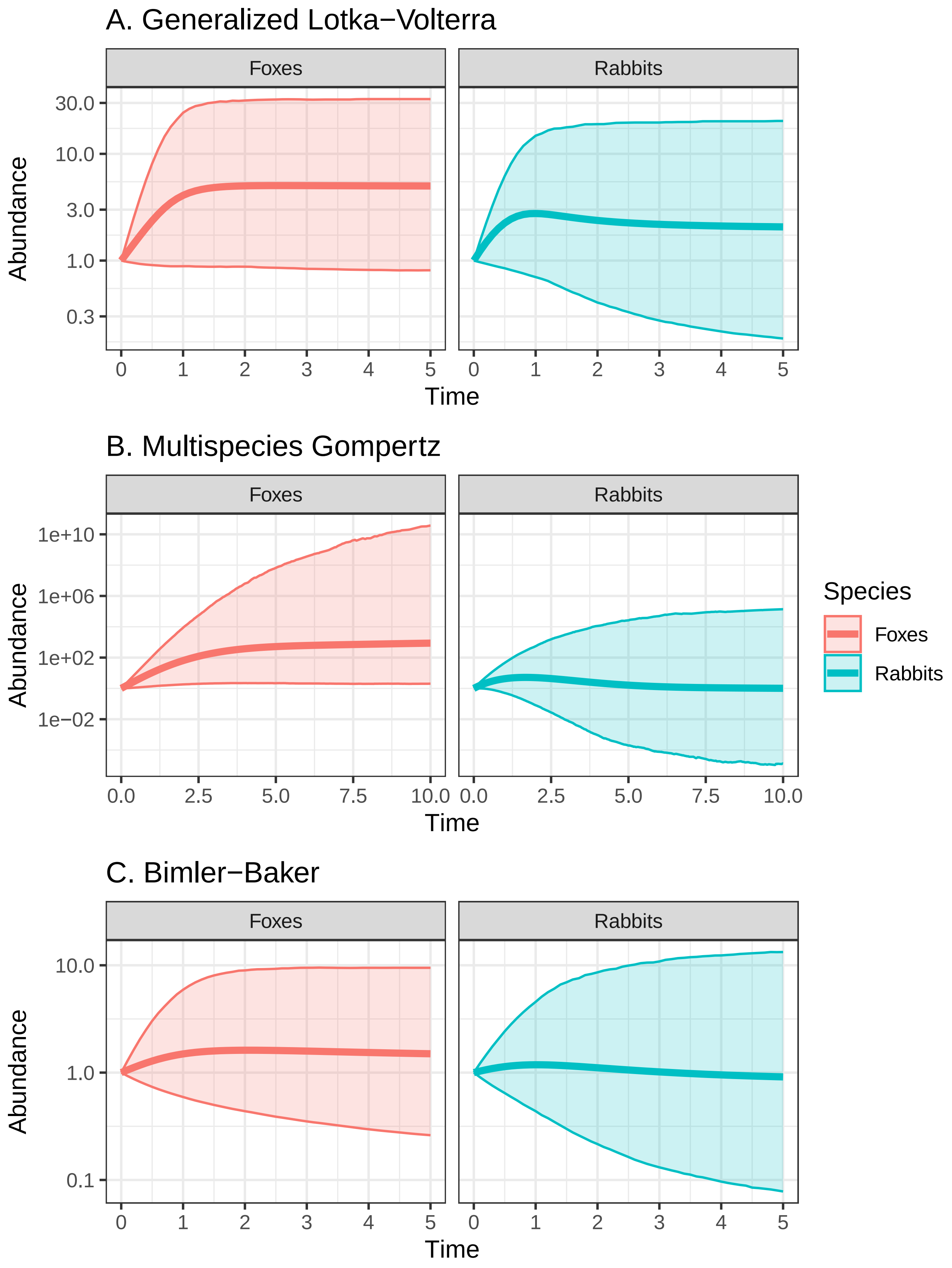


Figure S2: Example of abundances' predictions using the `plot_projections` function, for the three ecosystem models supported by the EEMtoolbox R-package. Thick lines represent the median abundance and thin lines bound the 95% prediction intervals.

##### B.4 Generating projections of future abundances of species scaled to the steady state

The function plot_projections can also be used to predict abundances scaled with the steady state, by setting the argument scaled to TRUE. In this case, the vector of initial abundances (initial_condition) is scaled to the steady state. In the following example, the vector c(0.9,1.1) signifies that the initial fox abundance is 0.9 of the steady state abundance, and the rabbit abundance is 1.1 the steady state abundance.

```
plot_projections(parameters_GLV, initial_condition = c(0.9,1.1), t_window = c(0,5), scaled=TRUE)
plot_projections(parameters_Gompertz, model="Gompertz",
                 initial_condition = c(0.9,1.1), t_window = c(0,5), scaled=TRUE)
plot_projections(parameters_Baker, model="Bimler-Baker",
                 initial_condition = c(0.9,1.1), t_window = c(0,5), scaled=TRUE)
```

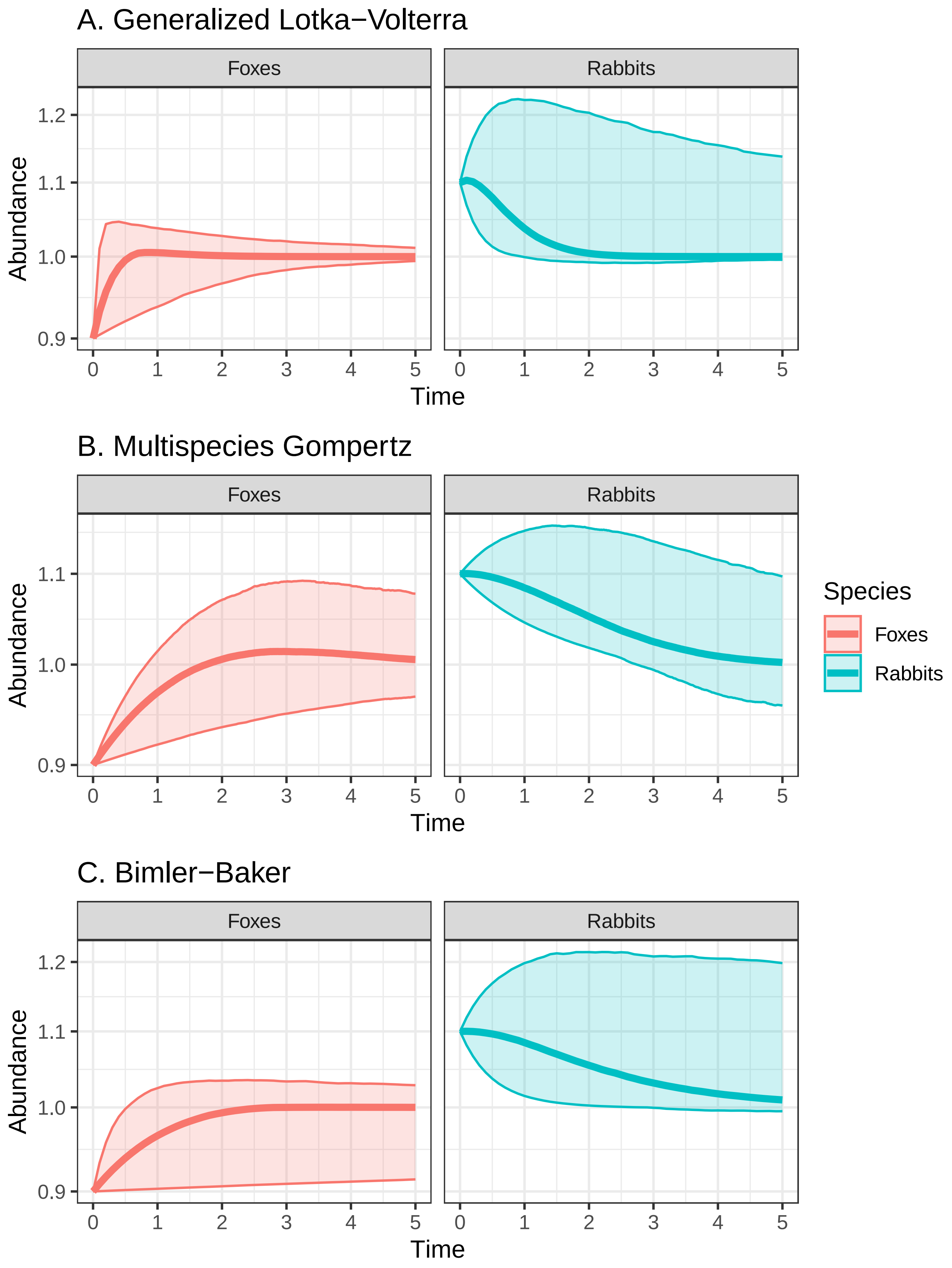


Figure S3: Example of abundances' predictions scaled to the steady state using the `plot_projections` function, for the three ecosystem models supported by the EEMtoolbox R-package. Thick lines represent the median abundance and thin lines bound the 95% prediction intervals. Here, the initial abundance of foxes is set to 90% of the steady state, and the initial abundance of rabbits to 110% of the steady state.

##### B.5 Standard EEM and SMC-EEM comparison

To demonstrate that the standard-EEM and SMC-EEM approaches give equivalent predictions (but at different computational costs), we generated 20,000 parameter sets with the standard EEM and the SMC-EEM approaches for the fox-rabbit example, and compare the abundances' predictions (Figure S4). For both species, the abundances predicted by parameters sampled with both methods overlap almost identically. This result suggests that SMC-EEM approach produces the same distribution of parameter sets as the standard-EEM approach, and substantial additional evidence for this point is detailed in (Vollert et al., 2024b). Any discrepancies are due to the stochasticity inherent in the random sampling processes used in both methods, often referred to as Monte Carlo error, and the finite number of parameter sets being generated. Generating these 20,000 parameter sets with the standard-EEM approach required 20 seconds, while the SMC-EEM approach required 35 seconds, both being computed on a Dell Laptop 11th Gen Intel(R) Core(TM) i7-1185G7 @ 3.00GHz, with 16 GB of RAM, 4 cores and 8 threads. Both sampling processes were parallelised in the code using the optional argument `"7L"` for `"n_cores"` in the package's `EEM` function (see Table 2), so that we were using 7 of the 8 available threads. However, it is important to note that the SMC-EEM approach will be drastically faster than the standard EEM approach for larger networks (Vollert et al., 2024b).

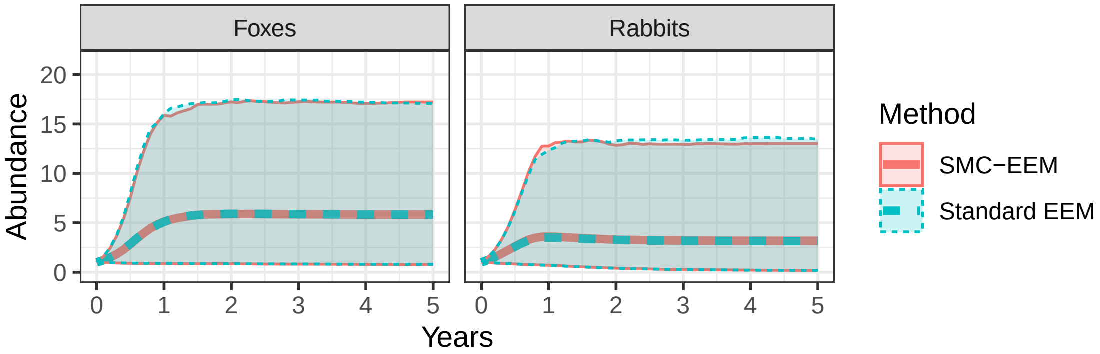


Figure S4: Example of abundances' predictions from parameter sets obtained with the standard accept-reject EEM approach and the SMC-ABC approach. Thick lines represent the median abundance and thin lines bound the 95% prediction intervals.

#### C Customized models

Our package is flexible to other ecosystem models customized by the user. Adapting the `EEM` function to a new ecosystem model requires defining a function to identify the unknown parameters of the model that need to be sampled (see function `args_function`), a function to summarise the feasibility and stability features of the new model (i.e. the equilibrium points and the real part of the Jacobian eigenvalues, see the function `summarise_ecosystem_features`) and a function to reconstruct the model from the sampled parameters (see `reconstruct_matrix_growthrates`).

Forecasting species abundances using the `plot_projections` function would require defining a function that returns the derivative of the model (see `derivative_func`).

### D Sihek case study

Table S1: Summary of interactions between species for the ecosystem network of the sihek case study (interaction matrix) and maximum growth rate for each species (Canessa et al., 2022). Coefficients of -1, 0 and 1 indicate that the abundance of the species in the column has a negative, negligible and positive effect respectively on the abundance of the species in the row.

| | Sihek | Seabirds | Terrestrial crabs | Carnivorous crabs | Cane spiders | Geckos | Cockroaches | Terrestrial arthropods | Native trees | $r_{\max}$ ($yr^{-1}$) |
|---|---|---|---|---|---|---|---|---|---|---|
| Sihek | -1 | 1 | 1 | 1 | 1 | 1 | 1 | 1 | 0 | 1.1 |
| Seabirds | -1 | -1 | 0 | -1 | 0 | 0 | 0 | 1 | 1 | 1.1 |
| Terrestrial crabs | -1 | 0 | -1 | -1 | 0 | 0 | 0 | 1 | -1 | 1.5 |
| Carnivorous crabs | -1 | 1 | 1 | -1 | 0 | 0 | 1 | 1 | -1 | 1.5 |
| Cane spiders | -1 | 0 | 0 | 0 | -1 | 1 | 1 | 0 | 0 | 0.39 |
| Geckos | -1 | 0 | 0 | 0 | -1 | -1 | 0 | 0 | 0 | 0.49 |
| Cockroaches | -1 | 0 | 0 | -1 | -1 | 0 | -1 | 0 | 0 | 3 |
| Terrestrial arthropods | -1 | 0 | 0 | -1 | -1 | -1 | 0 | -1 | 0 | 3 |
| Native trees | 0 | 1 | -1 | -1 | 0 | 0 | 0 | 1 | -1 | 3 |

### E Additional supplementary figures

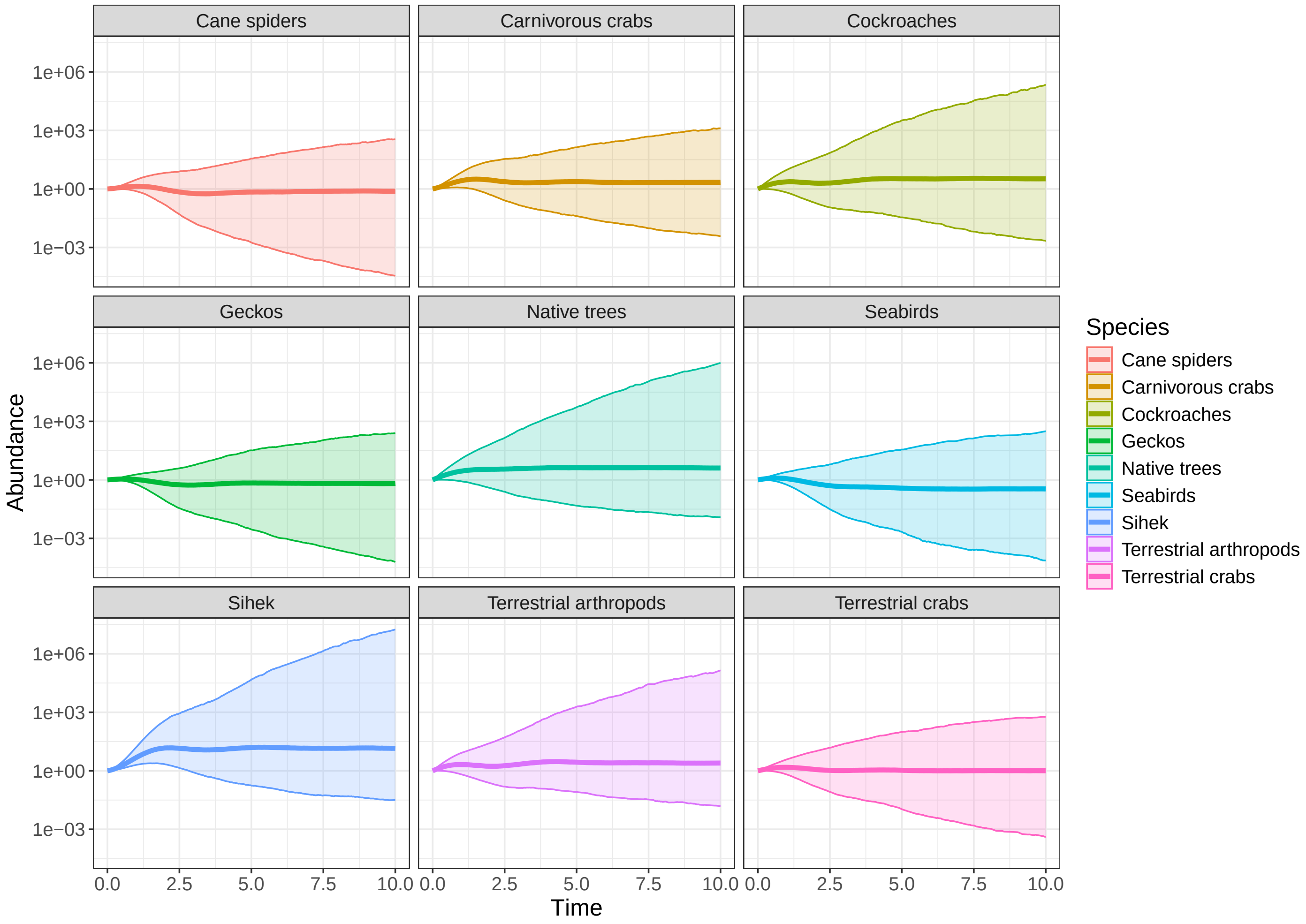


Figure S5: Forecasts of species abundances over 10 years for the sihek case study using the generalized Gompertz model. The initial abundance for each species is 1 individual per hectare. Thick lines represent the mean abundance and thin lines bound the 2.5% and 97.5 prediction intervals.

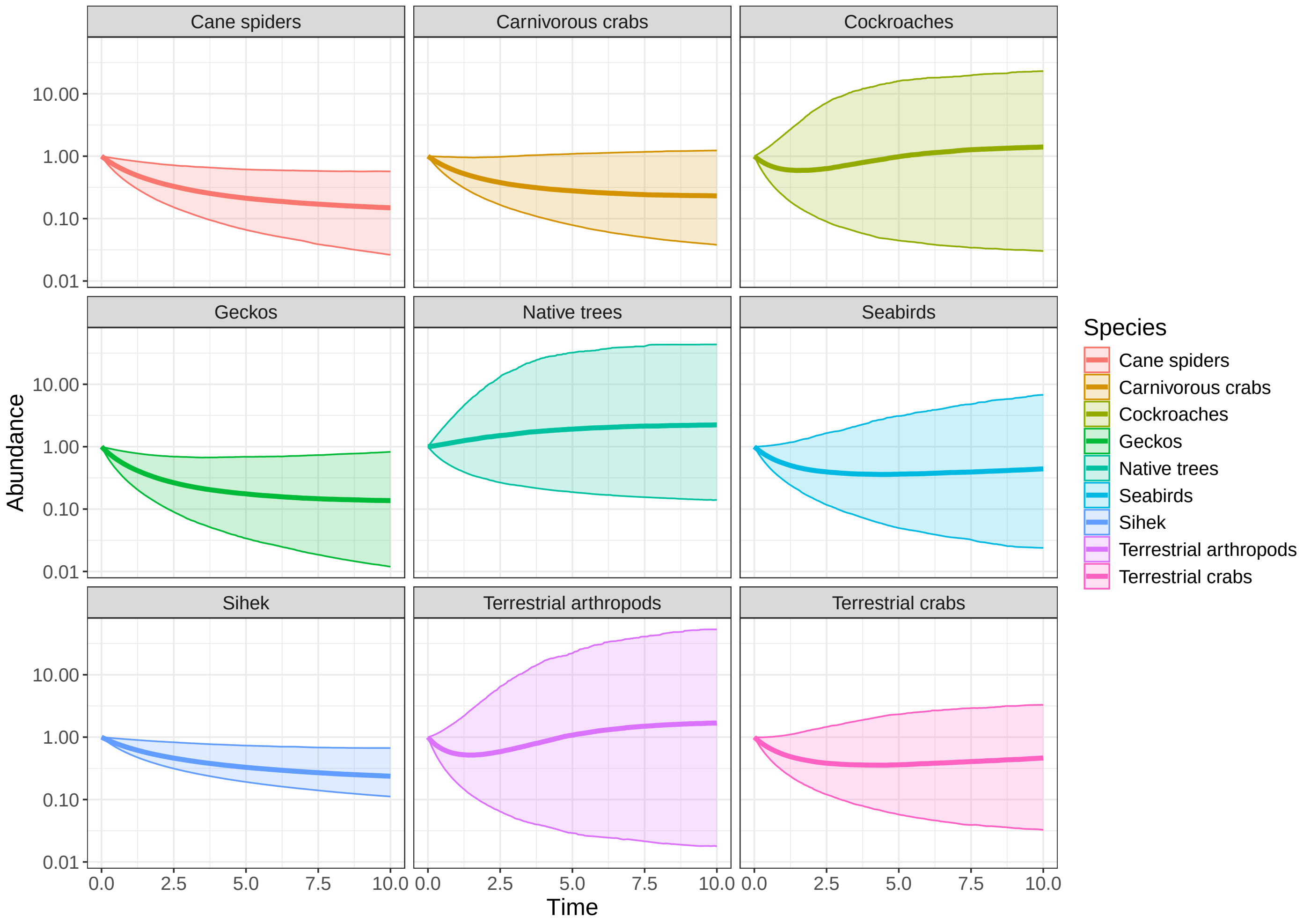


Figure S6: Forecasts of species abundances over 10 years for the sihek case study using the generalized Bimler-Baker model. The initial abundance for each species is 1 individual per hectare. Thick lines represent the mean abundance and thin lines bound the 2.5% and 97.5 prediction intervals.
